# Exploring Cellular Water Dynamics associated with Potassium Ion Changes Using Magnetic Resonance Imaging

**DOI:** 10.1101/2025.06.25.661446

**Authors:** Seong-Min Kim, Kyeongseon Min, Jung Seung Lee, Jang-Yeon Park

## Abstract

Potassium ions play a critical role in modulating cellular physiology, but their direct effects on water dynamics have not been fully explored. Here, we investigated how elevated potassium ion concentrations ([K⁺]) alter intracellular and extracellular water pools in comparison to hypoosmotic stress, using T2 and magnetization transfer (MT) parameters in a close-packed T-lymphocyte cell pellet model. Our findings reveal that the T2 increase primarily reflects an increase in intracellular free water concentration rather than a mere expansion of cell volume. Notably, [K⁺] elevation produced distinct cell swelling profiles and a smaller relative rise in free water at comparable volumetric changes compared to hypoosmotic stress, highlighting more complex mechanisms than straightforward osmotic effects. While T2 proved sensitive to shifts in intracellular water content, the bound pool increased linearly with cell volume expansion. These results underscore that [K⁺]-driven cell swelling diverges functionally from osmotic- driven cell swelling and demonstrate the viability of MRI-based approaches for probing K⁺-dependent cellular events.

## Introduction

Potassium ions play critical roles in cellular physiology, influencing numerous essential cellular processes, including signal transduction, volume regulation, and the maintenance of membrane potential^1–4^. Consequently, potassium ion modulation has been extensively employed in research aiming to elicit physiological responses and cellular alterations.

As cellular processes predominantly occur within an aqueous environment, physiological phenomena resulting from changes in potassium ion concentrations ([K^+^]) significantly affect water dynamics throughout biological systems. Water dynamics in biological contexts exhibit distinct states characterized by unique molecular behaviors and interactions^5–7^. Specifically, water can be categorized into dynamically slow interfacial water (or hydration layer), closely associated with macromolecular surfaces, and dynamically fast bulk-like free water, occupying most of the available intracellular and extracellular spaces^8–12^. Because both dynamically fast and slow water pools, together with the macromolecular components of cells, can undergo significant changes in a variety of physiological contexts^10,13,14^, elucidating their linked changes could shed light on how cells work and provide novel contrastive methods for spotting specific physiological events.

Recent studies using methods such as Raman spectroscopy, second harmonic generation (SHG), nuclear magnetic resonance (NMR), fluorescence microscopy, and fluorescence spectroscopy have demonstrated that these distinct water pools and macromolecules respond differently to various physiological stimuli^14–18^. Notably, SHG studies have reported measurable signal changes due to the reorientation of interfacial water molecules induced by alterations in membrane potential^19,20^. Since membrane potential variations are frequently associated with changes in [K^+^], these findings highlight that [K^+^]-related physiological phenomena, such as membrane potential alterations, can markedly impact water dynamics at cellular interfaces.

Among non-invasive imaging techniques, magnetic resonance imaging (MRI) - particularly quantitative magnetization transfer (MT) imaging - can effectively distinguish between free water pool and bound pool by examining the energy transfer dynamics between these pools^21,22^. Although MRI does not provide as high spatial resolution as optical microscopy, its non-invasive and wide-ranging imaging capabilities offer substantial practical advantages. MRI-based approaches thus hold considerable potential for monitoring physiological conditions through measurements of water dynamics, potentially providing novel insights into underlying cellular mechanisms *in vivo*.

Previously, our group reported a measurable increase in the spin-spin interaction relaxation time (T2) and a decrease in one of the MT parameters, the pool size ratio (PSR, the ratio of bound to free water pools), in response to increasing [K^+^] in both *in vitro* and *in vivo* conditions, demonstrating that these MRI parameters are sensitive to changes in cellular physiology^23^. Other studies demonstrated that the increase in T2 associated with cell swelling occurs from dilution effects^24–26^, which may also be interpreted in terms of their potential link to increased intracellular free water content. Additionally, some MT studies mentioned that the bound pool includes hydration layers and macromolecules^21,27,28^.

In this study, we explored water dynamics in cellular processes associated with [K^+^] changes using MRI. In particular, we attempted to explain the changes in T2 relaxation time and MT parameters in MRI induced by [K^+^] changes in terms of cellular water dynamics based on intracellular structural modeling *in vitro*. In addition, we also investigated whether cell swelling due to [K^+^] changes is the sole factor contributing to the changes in these MRI parameters, compared to cell swelling induced by changes in osmolarity as a control condition, and examined whether there were differences in water dynamics between [K+] elevation and hypoosmotic stress conditions.

## Results

### One-dimensional (1D) cell pellet imaging & mean cell volume measurements

We used T-lymphocyte cells (or Jurkat) in pellet form, where the cells were modeled as close-packed spherical structures. To ensure stable cell volumes following cell volume regulation, sufficient stabilization time was allowed without compromising cell viability. Suspended cells were loaded into the perforated wells of an appropriately shaped acrylic phantom (Figure 1a). To induce cell swelling, cells were suspended in media prepared under hypoosmotic stress and [K^+^] elevation for an hour. For hypoosmotic stress, additional osmotic concentrations of -10 mM, -20 mM, and -30 mM were added to the control media; for [K^+^] elevation, [K^+^] of 20 mM, 40 mM, and 80 mM were used.

**Figure 1.**
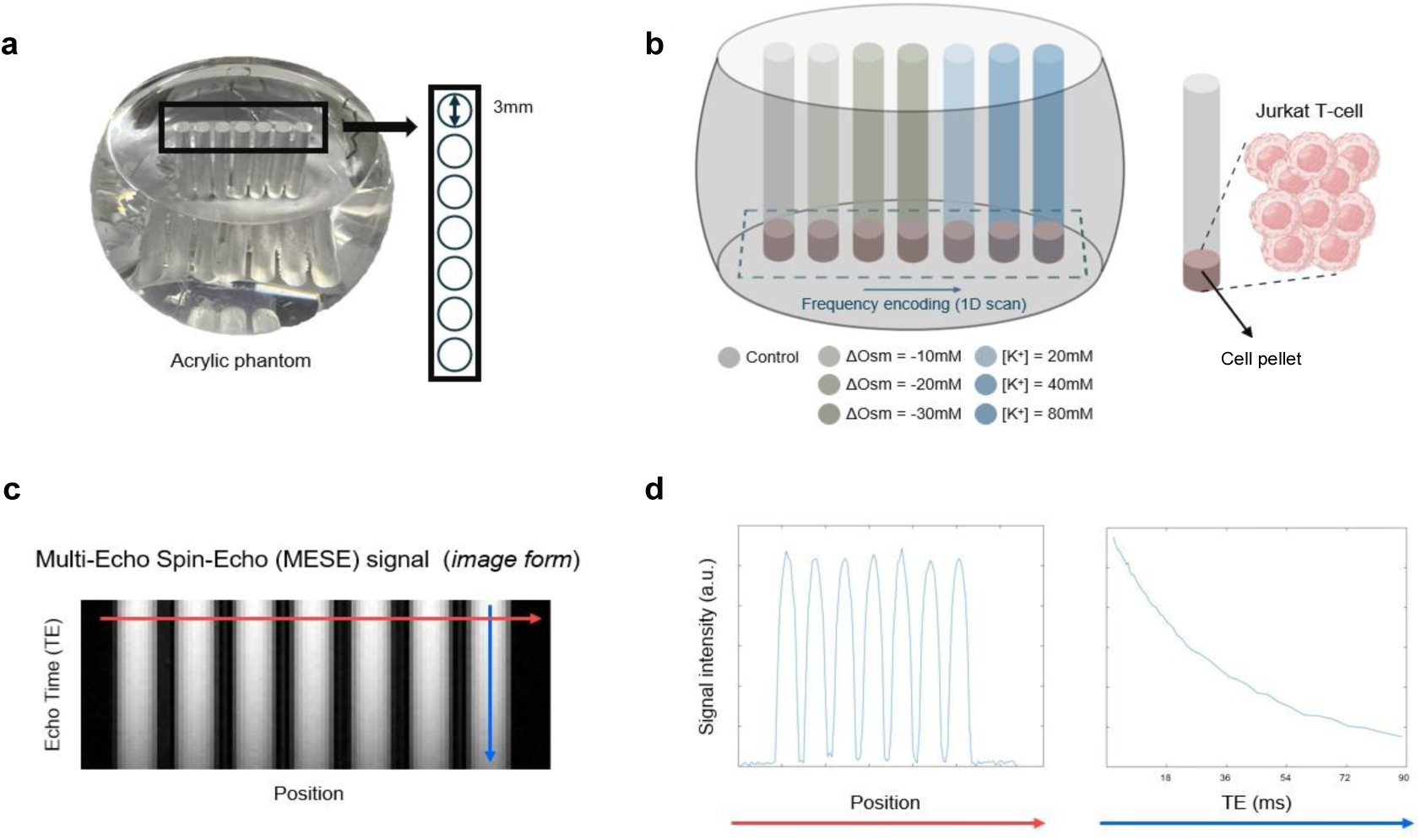
Acrylic phantom containing cells and MRI signals. **a.** Photo of a fabricated acrylic phantom with 7 wells. **b.** Illustration of the acrylic phantom containing cells in pellet form. Each well of the acrylic phantom contains T-lymphocyte cells (Jurkat) suspended under different conditions. Centrifugation was applied to pellet the cells. **c.** A series of 1D MR images for various echo times (TEs) with respect to position. **d.** Signal intensity profiles along the red solid line (left) and blue solid line (right) shown in **c**.

Cells were centrifuged to form cell pellets, which were then prepared to a sufficient thickness (> 1 mm) after accumulating under different media conditions (Figure 1b). 1D imaging was performed to estimate MT and T2 across cell pellets in these different media. A multi-echo spin-echo sequence with variable echo times (TE) was used for T2 estimation, and Figure 1c shows the 1D images for different TEs. In Figure 1d, an example of 1D signal profile of cell pellets in media with different conditions, contained in separate wells, is shown in the position dimension (along the red solid arrow line in Figure 1c), together with the signal intensities in the TE dimension (along the blue solid arrow line in Figure 1c).

To estimate cell volume expansion, microscopy images were acquired as shown in Figure 2a. Assuming a spherical cell shape, cell volume was calculated based on the measured cell radius. Figure 2b shows the cell volume distribution obtained by measuring the volume for each of 1,000 cells. These results demonstrate that both media prepared for elevated [K^+^] and hypoosmotic stress conditions induced significant levels of cell swelling, leading to an expansion of the average cell volume (〈*V*_*c*_〉).

**Figure 2.**
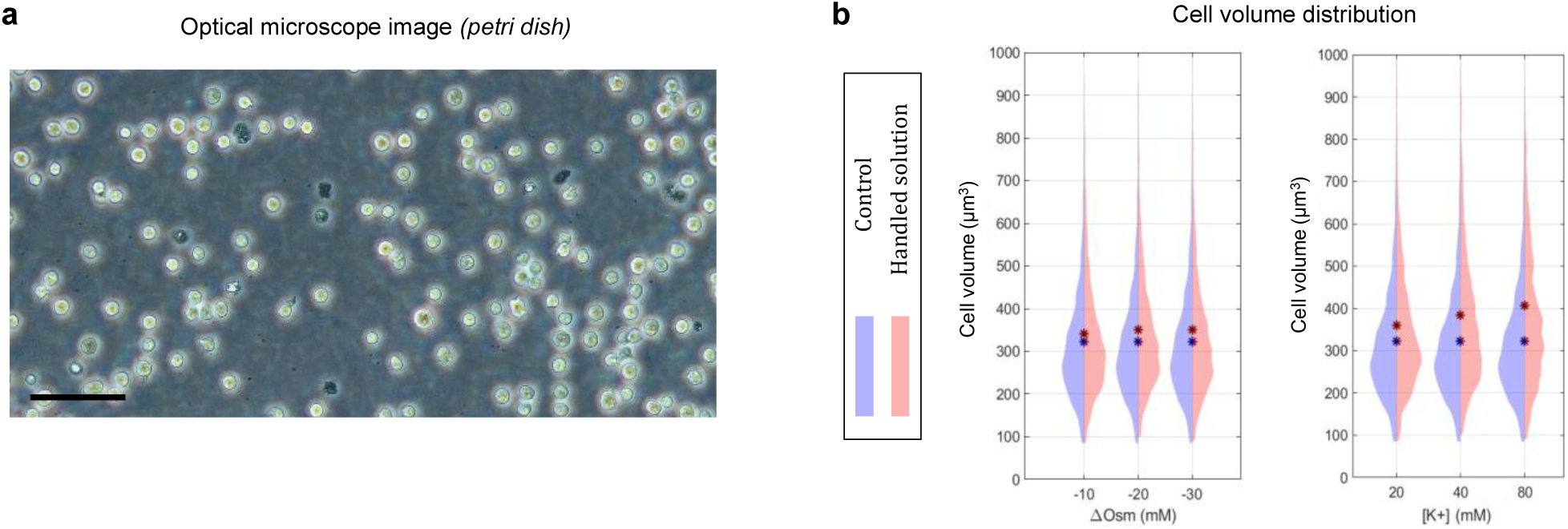
Microscope image and cell volume distribution. **a.** Microscope image of the cells acquired at 20x magnification. The size of the scale bar at bottom left is 50 μm. **b.** Cell volume distribution displayed using a violin plot. The radius of each cell cross-section was measured in **a** to estimate the cell volume distribution.

### Changes in free water pool by cell swelling

To understand cell water dynamics during cell swelling, MT parameters were measured using an inverse recovery multi-echo spin echo sequence. Free water and bound pools were expressed in terms of longitudinal magnetization at equilibrium state (*M*_*f*,∞_ and *M*_𝑏,∞_, respectively, defined in Eqs. [1] and [4] in Materials and Methods). To better understand what the estimated *M*_*f*,∞_ represents in cell pellet conditions, we first modeled the cell pellet as a close-packed pellet structure as shown in Figure 3a. In this model, when cells expand, their geometric configuration with adjacent cells changes, reducing the number of cells per unit volume (cell density, *ρ*_*cell*_). In addition, if the unit volume is sufficiently larger than the size of the cell expansion, the ratio of intracellular space per unit volume (*v*_*i*_) and the ratio of extracellular space per unit volume (1−*v*_*i*_) remain constant regardless of changes in the cell volume^29,30^. Figure 3a illustrates the increase in average cell volume 〈*V*_*c*_〉 and the decrease in cell density *ρ*_*cell*_ during cell swelling in a three-dimensional (3D) structure of unit volume. Figure 3b illustrates the two-dimensional (2D) cross section of Figure 3a, including the organelles (orange) inside the cell, before and after cell expansion. The yellow square box represents the boundary of the unit volume cross-section. To better illustrate the maintenance of *v*_*i*_ (or 1−*v*_*i*_) during cell swelling, Figure 3c combines Figures 3a and b, representing the spherical cell volume as a rectangular solid.

**Figure 3.**
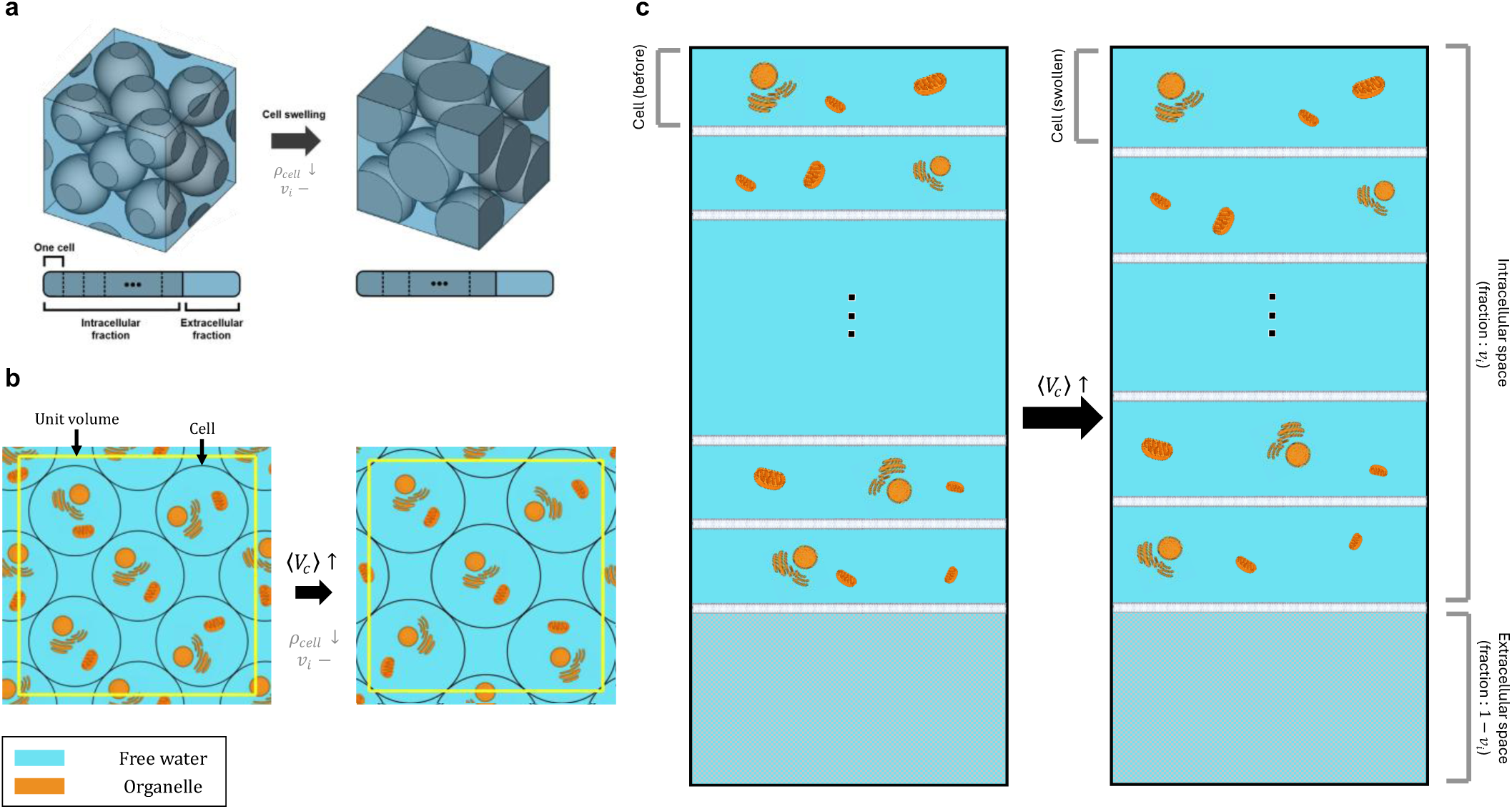
Illustration of a close-packed model of cell pellet before (left) and after cell swelling (right). **a.** Close-packed spherical cell structure of unit volume. **b.** 2D cross-section including organelles of the 3D spherical cell structure. The yellow square box represents the boundary of the unit volume cross-section in **a**. It is well shown that after cell swelling, cell density decreases and the density of subcellular structures, including organelles and cell membranes, also decreases. **c.** Combination of **a** and **b** representing the spherical cell volume as a rectangular solid. While the number of cells per unit volume decreases, the intracellular and extracellular space fractions remain constant.

As shown in Figure 3, cell swelling leads not only to a decrease in *ρ*_*cell*_, but also to a decrease in the proportion of subcellular structures, including organelles and cell membranes, per unit volume. In this case, the intracellular space expands due to the influx of free water, and the concentration of intracellular free water pool also increases due to the fixed ratio of *v*_*i*_. This situation corresponds to a phenomenon called ‘dilution’ in previous studies reporting an increase in T2 during cell swelling^25,26^.

On the other hand, for the extracellular space, since the media used to induce cell swelling in this study differ only in their ionic concentrations on the millimolar scale, it can be assumed that such minor variations would not significantly alter the concentration (or ^1^H proton density) of the extracellular free water pool under different conditions. This assumption allows us to ignore concentration changes in the extracellular free water pool. Therefore, given that *v*_*i*_ (or 1−*v*_*i*_) remains constant in the close-packed cell pellet model during cell swelling, the change in *M*_*f*,∞_ can be best explained by the change in the intracellular free water concentration.

Figure 4a shows *M*_*f*,∞_ evaluated from the data measured in the two conditions used to induce cell swelling, i.e., osmotic changes (ΔOsm) and [K^+^] changes, respectively. A significant increase in *M*_*f*,∞_was observed in both cases of hypoosmotic stress and [K^+^] elevation. In addition, as shown in Figure 4b, cell swelling under the two conditions revealed different relationships between *M*_*f*,∞_ and 〈*V*_*c*_〉. Under hypoosmotic stress, the cell volume became saturated beyond a certain threshold, whereas *M*_*f*,∞_ increased linearly. These observations can be understood from a cell physiology perspective. In other words, under hypotonic conditions, cells initially swell, but eventually partially recover their size over time through a process called regulatory volume decrease (RVD). Therefore, under high hypoosmotic stress for a sufficient period of time, cells ultimately maintain their volume due to RVD regardless of the degree of stress, whereas the intracellular free water pool concentration adjusts linearly to the external osmotic environment to maintain osmotic balance^33^. In contrast, as [K^+^] increased, both cell volume and intracellular free water pool concentration increased linearly. These results may be because, unlike the hypotonic environment mentioned above, high [K^+^] conditions do not induce a decrease in cell volume after swelling^1,32,34,35^. As with other ions and osmolytes, the influx or efflux of K^+^ is accompanied by the influx or efflux of free water molecules, as the cell volume expands or contracts, respectively. In summary, these findings demonstrate that two different mechanisms, [K^+^] changes and osmotic changes, have distinct trends in cell volume expansion and free water pool concentration changes, which can be measured and analyzed by MRI.

**Figure 4.**
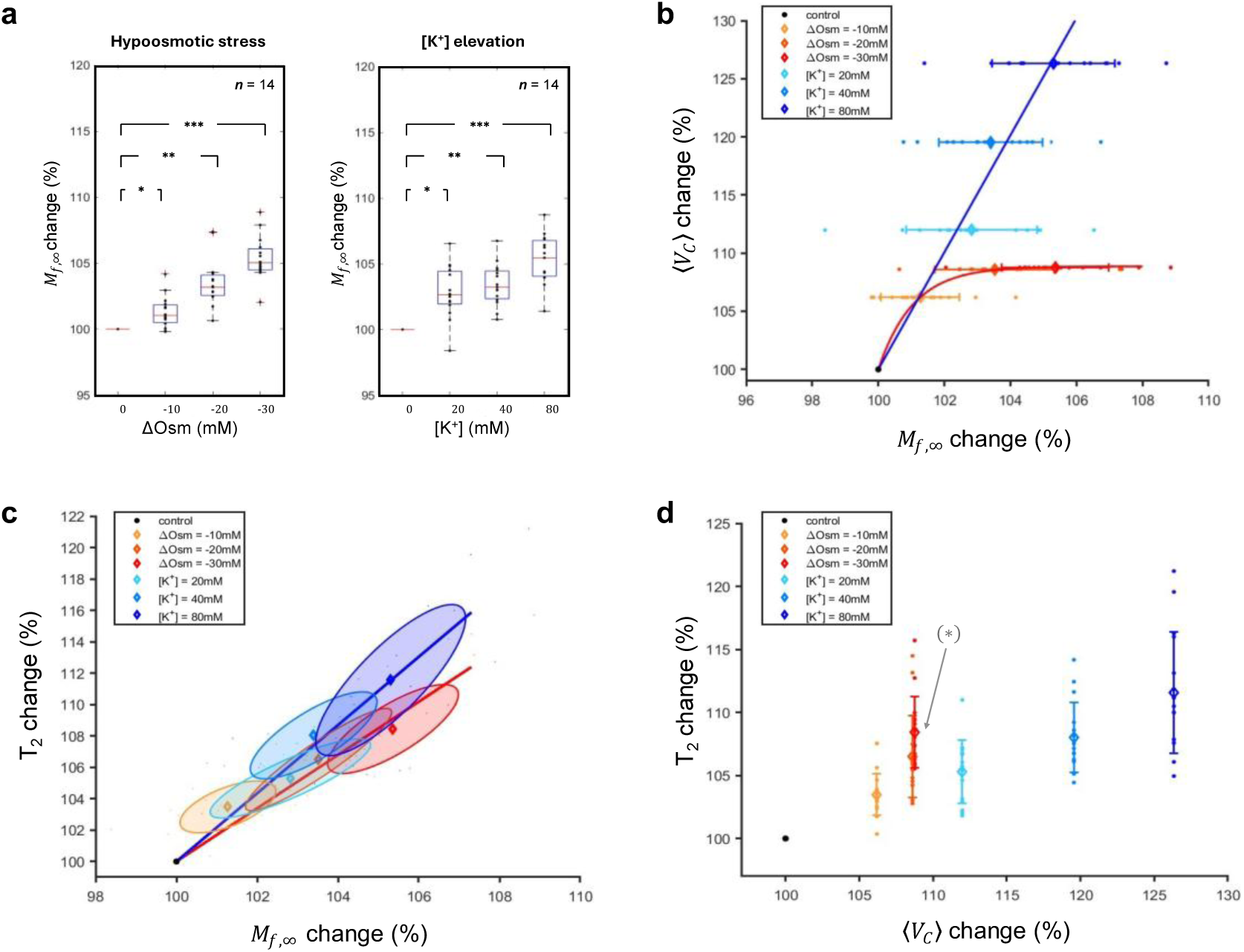
Estimation of free-water pool changes based on quantitative MT and modeling. a. *M*_*f*,∞_ changes under hypoosmotic stress (left) and [K^+^] elevation (right) conditions (n 14 scans). *M*_*f*,∞_ increased with statistical significance in both cases. b. 〈*V*_*c*_ 〉 changes with respect to *M*_*f*,∞_ changes. 〈*V*_*c*_ 〉 increased as *M*_*f*,∞_ increased, but the two conditions showed different trends. Under hypoosmotic stress, 〈*V*_*c*_〉 was saturated beyond a certain threshold, but increased linearly under [K^+^] elevation. **c**. T2 changes with respect to *M*_*f*,∞_changes. In both cases, T2 increased linearly, but with a steeper slope under [K^+^] elevation. The shaded ellipses represent the distribution of data points, with the major and minor axes corresponding to the variance obtained from the principal component analysis (PCA) of each data point. **d**. T2 changes with respect to 〈*V*_*c*_〉 changes. T2 increased linearly with increasing 〈*V*_*c*_〉, but under hypoosmotic stress conditions of -20mM and -30mM, a statistically significant difference in T2 was observed despite indistinguishable cell volume increase. Statistical significance was marked with asterisks (*: p < 0.05; **: p < 0.01; ***: p < 0.001). (*M*_*f*,∞_: the longitudinal magnetization of free water pool at equilibrium state, 〈*V*_*c*_ 〉: average cell volume).

Figure 4c shows that T2 increased linearly with increasing *M*_*f*,∞_ in both cases of hypoosmotic stress and [K⁺] elevation, suggesting that the rise in intracellular free water concentration contributed to this T2 increase irrespective of the cell swelling mechanism. If the increase in free water within the intracellular space is regarded as intracellular dilution, this finding is consistent with previous studies.^25,26^. The contribution of bound pool to T2 increase may be negligible because bound pool is very localized and T2 is significantly shorter (< 1 ms) than the TE used here (> 8 ms). On the other hand, for the same level of *M*_*f*,∞_ increase, elevation of [K+] resulted in a greater increase in T2 than hypoosmotic stress, indicating that the two cell-swelling mechanisms can lead to different T2 changes even at similar levels of intracellular dilution. Since increasing [K^+^] resulted in a larger volume at comparable *M*_*f*,∞_levels (Figure 4b), the higher T2 when increasing [K^+^] may be due to the relatively larger intracellular space per cell than when decreasing osmotic pressure (or under hypoosmotic stress). According to previous studies, T2 varies with the size of spatial restriction^36–39^, that is, in more open and spacious intracellular environments, T2 is larger than in dense and confined spaces. In summary, T2 changes differed by two different mechanisms, [K^+^] changes and osmotic changes, and cell swelling due to increased [K^+^] resulted in a larger T2 increase than cell swelling due to decreased osmotic pressure.

Figure 4d shows how T2 changes vary with 〈*V*_*c*_〉. At the same 〈*V*_*c*_〉, the T2 increase under hypoosmotic stress was greater than that under [K^+^] elevation. From a cellular physiological perspective, this may be explained by RVD causing greater intracellular dilution (or higher intracellular free water concentration) under hypoosmotic stress, leading to an increase in T2. In particular, when ΔOsm -20 mM or -30 mM, there was a significant difference in T2 increase even with similar 〈*V*_*c*_〉, suggesting that T2 can increase by the dilution effect without cell volume expansion.

Taken together, the free water pool increased during cell swelling, but the relationship between an increase in *M*_*f*,∞_ and cell expansion was determined by the mechanisms of cell swelling (Figure 4b). In both mechanisms of hypoosmotic stress and [K⁺] elevation, the change in T2 depended linearly on the change in *M*_*f*,∞_, the slope of which depended on physiological factors such as cell volume and the associated intracellular dilution.

### Changes in bound pool per cell

As shown in Figure 5a, the ratio of bound to free water pools, PSR, decreased in response to both hypoosmotic stress and [K^+^] elevation, consistent with our previous reports^23^. Using PSR and *M*_*f*,∞_, the longitudinal magnetization of bound pool at equilibrium state, *M*_𝑏,∞_, was also calculated (Figure 5b, for detailed calculations, see the data analysis of Materials and Methods). No significant change in *M*_𝑏,∞_was observed due to cell swelling under hypoosmotic stress. In contrast, for [K^+^] elevation, a statistically significant decrease in *M*_𝑏,∞_(p < 0.05) was observed when [K^+^] was 40 mM and 80 mM. These results suggest that the decrease in PSR under hypoosmotic stress mainly results from the increase in *M*_*f*,∞_(Figure 4a), whereas the decrease in PSR with increasing [K^+^] is due to both the increase in *M*_*f*,∞_(Figure 4a) and the decrease in *M*_𝑏,∞_.

**Figure 5.**
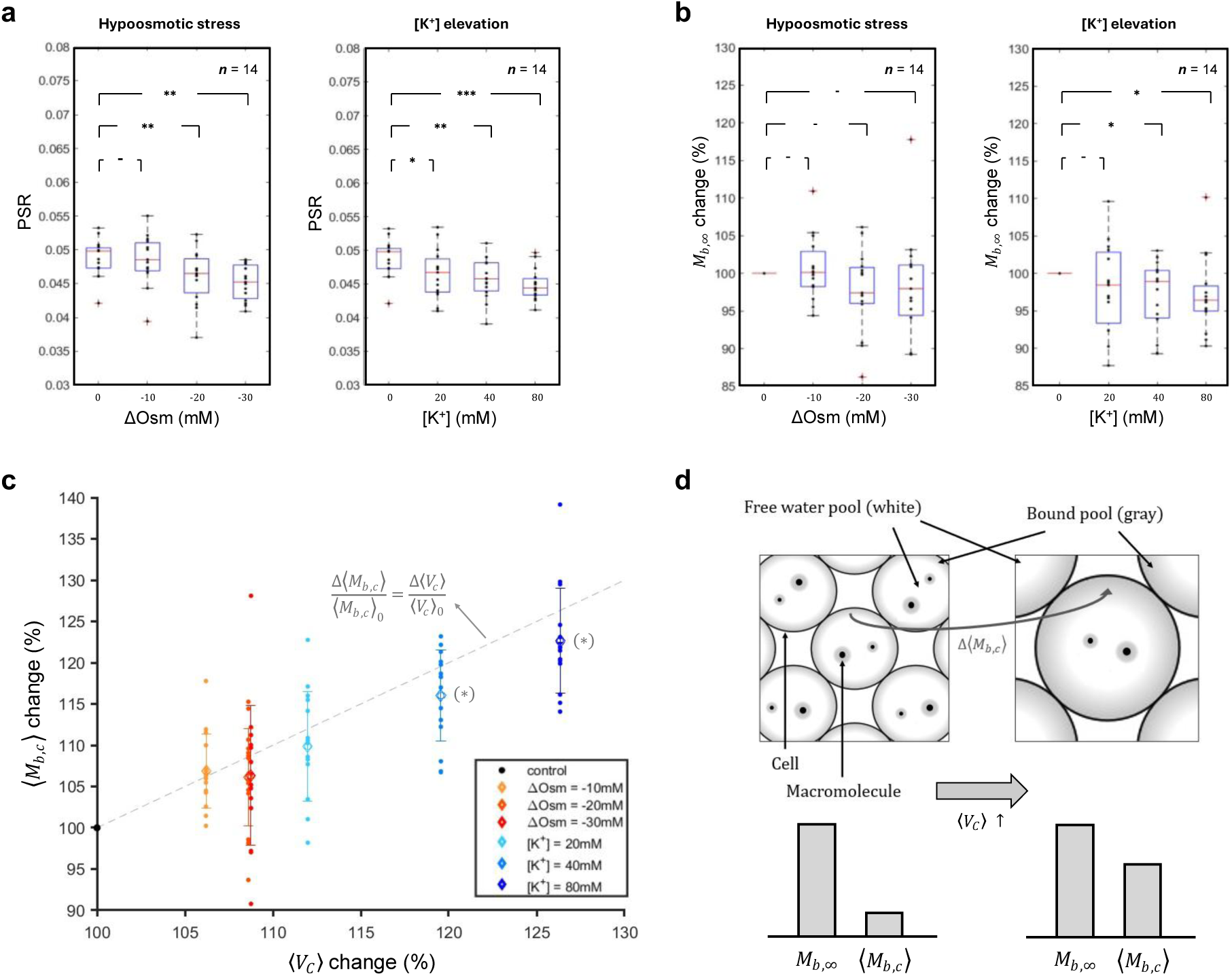
Estimation of bound pool changes based on quantitative MT and modeling. **a.** PSR changes under hypoosmotic stress (left) and [K^+^] elevation (right) conditions (n 14 scans). PSR decreased with statistical significance in both cases. **b.** *M*_𝑏,∞_ changes under hypoosmotic stress (left) and [K^+^] elevation (right) conditions (n 14 scans). *M*_*f*,∞_was calculated using PSR and *M*_*f*,∞_. Unlike the osmotic stress condition, [K+] elevation showed a statistically significant reduction in *M*_𝑏,∞_ at high [K+], but under hypoosmotic stress, no statistically significant decrease was observed. **c.** 〈*M*_𝑏,*c*_〉 changes with respect to 〈*V*_*c*_ 〉 . 〈*M*_𝑏,*c*_〉 was calculated based on a close-packed cell pellet model. The gray dashed line represents the trend of increasing 〈*M*_𝑏,*c*_〉 proportionally with cell volume expansion. In cases where *M*_𝑏,∞_ decreased, the data points deviated from the gray dashed line. **d.** Illustration of the close-packed spherical cell pellet model including free water and bound pools. While the total *M*_𝑏,∞_ within a voxel remains constant, 〈*M*_𝑏,*c*_〉 increases as the cell volume expands. Statistical significance was marked with asterisks (-: p > 0.05; *: p < 0.05; **: p < 0.01; ***: p < 0.001). (*M*_𝑏,∞_: the longitudinal magnetization of bound pool at equilibrium state, 〈*M*_𝑏,*c*_〉: the average longitudinal magnetization of bound pool per cell).

On the other hand, since changes in *M*_𝑏,∞_, which represents the bound pool within a voxel, do not directly reflect changes in the bound pool of individual cells, we also estimated the average longitudinal magnetization of the bound pool per individual cell, 〈*M*_𝑏,*c*_〉, which is calculated by *M*_𝑏,∞_/*ρ*_*cell*_ (see Eq. [5] in Materials and Methods). Figure 5c shows the increase in 〈*M*_𝑏,*c*_〉 according to the relative change in 〈*V*_*c*_〉 under two conditions of hypoosmotic stress and [K^+^] elevation, respectively. The increase in 〈*M*_𝑏,*c*_〉 is expected from the results that the decrease in *M*_𝑏,∞_ during cell swelling was about 10 even at maximum, e.g., under high [K+] conditions (Figure 5b), whereas the increase in 〈*V*_*c*_〉 was about 25 (Figure 5c) and thus the decrease in *ρ*_*cell*_was about 25 under the same conditions. The gray dashed line in Figure 3c represents a straight line along which the relative change in 〈*M*_𝑏,*c*_〉 with respect to the initial 〈*M*_𝑏,*c*_〉_0_ before cell swelling, Δ〈*M*_𝑏,*c*_〉/〈*M*_𝑏,*c*_〉_0_, is equal to the relative change in 〈*V*_*c*_〉 with respect to the initial 〈*V*_*c*_〉_0_ before cell swelling, (Δ〈*V*_*c*_〉/〈*V*_*c*_〉_0_). Hypoosmotic stress conditions where no statistically significant changes in *M*_𝑏,∞_were observed corresponded to trends in this gray line. The conditions for increasing [K^+^] also showed a similar trend overall, but when [K^+^] was 40 mM and 80 mM, this trend deviated and the change in 〈*M*_𝑏,*c*_〉 became smaller due to the decrease in *M*_𝑏,∞_(Figure 5b) and tended to be located below the gray line. These results demonstrate that during cell swelling, the bound pool increases proportionally with cell volume expansion and that additional factors appear to contribute to the decrease in the bound pool in high [K^+^] environments. Figure 5d illustrates how 〈*M*_𝑏,*c*_〉 grows in a close-packed cell pellet model as 〈*V*_*c*_〉 expands and *ρ*_*cell*_ decreases accordingly.

## Discussion

In this study, we investigated water dynamics associated with cell swelling under two conditions: [K^+^] elevation and hypoosmotic stress in a close-packed cell pellet model, using MT parameters and T2 relaxation time in MRI. In particular, we investigated whether the changes in these MRI parameters due to [K^+^] elevation are solely due to cell swelling from a mechanical point of view.

According to our results, cell swelling in both conditions induced measurable increases in T2, which were linearly correlated with increases in intracellular free water concentration (or dilution) rather than cell volume per se, consistent with previous studies^24–26^. Interestingly, for the same change in cell volume (〈*V*_*c*_〉), hypoosmotic stress conditions resulted in higher T2 and larger free water pool (*M*_*f*,∞_) than elevated [K^+^] conditions. In addition, cell swelling due to hypoosmotic stress exhibited cell volume saturation due to RVD, whereas intracellular free water concentration continued to increase linearly with increasing hypoosmotic stress. In contrast, increasing [K^+^] induced larger cell volume expansion at the same dilution level compared to hypoosmotic stress, suggesting that cell volume responses to dilution vary depending on the relevant physiological mechanisms.

Our results also demonstrated that the PSR decreased significantly in response to both hypoosmotic stress and [K^+^] elevation, aligning with our previous findings^40^. This decrease in PSR coincided with a statistically significant increase in *M*_*f*,∞_ under both conditions. The bound pool (*M*_𝑏,∞_) estimated by PSR and *M*_*f*,∞_ showed no statistically significant changes under hypoosmotic stress conditions, but showed statistically significant decreases at higher [K^+^] (e.g., 40 mM and 80 mM). The relatively larger change in *M*_*f*,∞_ compared to *M*_𝑏,∞_ indicates that the decrease in PSR primarily reflects an increase in the free water pool rather than a reduction in the bound pool.

Examining the change in bound pool per cell (〈*M*_𝑏,*c*_〉), calculated based on a close-packed cell pellet model, we observed an increase in 〈*M*_𝑏,*c*_〉 proportional to cell swelling, which can be interpreted as an adaptive response to the expansion of cell volume, including *de novo* synthesis of structural proteins, expansion of cell membrane surface area, and consequent growth of interfacial water. A recent study by Watson et al. showed that cells buffer changes in water potential by unfolding intrinsically disordered protein domains, thereby increasing protein surface area and thus expanding the associated slow-exchange hydration layer^13^. Our finding is also consistent with the increase in the slow diffusion pool (SDP) fraction reported in diffusion-functional MRI studies of neuron swelling^41–43^.

However, under high [K^+^] conditions, the increase in 〈*M*_𝑏,*c*_〉 during cell swelling was less pronounced compared to that induced by hypoosmotic stress. These differences are consistent with our observation that under hypoosmotic stress, cell volume expands only up to a certain amount due to the RVD response, whereas under [K⁺] elevation, it increases linearly with increasing [K⁺]. Furthermore, these results support that although both environments of [K⁺] elevation and hypoosmotic stress lead to an overall increase in cell volume, they involve different physiological processes in cell swelling.

As one possible physiological process uniquely related to changes in 〈*M*_𝑏,*c*_〉 under [K⁺] elevation, the change in membrane charge gradient due to the rise in [K^+^] could affect the dynamics of interfacial polar water molecules that are located near the membrane surface and likely constitute part of the bound pool alongside macromolecules. Previous studies using SHG demonstrated measurable signal changes resulting from reorientation of interfacial water molecules driven by membrane potential variations, including those caused by [K^+^] elevation^19,20,44,45^. Additionally, Raman spectroscopy studies have shown that reductions in surface electric potential correlate with decreased interfacial water signals, even in non-biological systems^46^. These studies agree with the results on water dynamics reported in this study. In other words, as a redistribution process, the interfacial water dissociated due to the change in membrane potential may move into the free water pool, thereby decreasing the bound water pool and increasing the signal of the free water pool. Interestingly, one of the quantitative MT parameters not presented above, the MT exchange rate 𝑘_𝑏*f*_, showed a statistically significant increase upon [K⁺] elevation, but no significant change was detected during hypoosmotic stress (Supplementary Fig. S1b). These differences in 𝑘_𝑏*f*_may be expected to serve as a potential source of MRI contrast to detect changes in membrane potential.

On the other hand, physiological processes that are not related to changes in membrane potential may also be involved. In other words, since K⁺ acts not only as volume regulating ions but also as a pleiotropic signaling factor, physiological processes associated with [K⁺] elevation might also involve other non-exclusive mechanisms, such as selective synthesis or redistribution of intracellular macromolecules required to tolerate sustained high-[K⁺] conditions, or alternative adaptation programs that replace the canonical RVD when cell volume increases beyond the osmotic set point. To clarify whether the differences in 〈*M*_𝑏,*c*_〉 between [K⁺] elevation and hypoosmotic stress conditions are primarily attributed to changes in membrane potential, it would be helpful to perform further studies that include MT studies using artificially controlled cell membranes or proteins whose surface charge can be manipulated without the intervention of complex cellular functions (e.g., homeostatic regulation).

Based on these findings, we hypothesize that the relatively smaller increase in 〈*M*_𝑏,*c*_〉 under [K^+^] elevation conditions could be due to a decrease in interfacial water caused by changes in membrane potential. This interpretation suggests that interfacial water, modulated by electrical properties of the membrane, may also play a role in the bound pool dynamics in [K^+^] elevation-induced cell swelling. To substantiate this hypothesis, future studies should explore the direct relationship between interfacial water and the bound pool measured in qMT experiments. Furthermore, investigating how changes in dynamically slow bound water induced by changes in membrane potential could be leveraged for enhanced contrast in MRI signals could potentially expand the application of MRI in studying cellular and membrane dynamics. For example, some MRI studies have practically applied changes in water dynamics in cases such as multiple sclerosis (MS) or cytotoxic edema^47–49^.

Finally, the estimated changes in 〈*M*_𝑏,*c*_〉 associated with [K+] elevation in this study may serve as a potential source of MRI contrast to detect nerve impulses that occur in milliseconds with negligible volume change and are accompanied by membrane potential changes during neuronal activity. For example, our simulations based only on the change in 〈*M*_𝑏,*c*_〉 at a fixed cell volume yielded a 1 increase in signal change when 6.9 of the bound pool was redistributed into the free water pool (Supplementary Fig. S2b). This may be because hydrogen protons (^1^H) in the bound pool, previously ‘invisible’ due to their very short T2, became detectable, effectively increasing the apparent proton density.

In conclusion, MT and T2 measurements using MRI revealed that cell swelling induced by either [K^+^] elevation or hypoosmotic stress led to an increase in the free water pool and T2, and that in this case, the T2 increase was primarily due to an increase in the intracellular free water pool (or dilution). In addition, for an equivalent increase of cell volume, cell swelling due to [K^+^] elevation resulted in a smaller increase in the bound pool per cell, whereas cell swelling due to hypoosmotic stress showed a larger increase in both T2 and *M*_*f*,∞_. These findings show that K^+^-induced cell swelling involves different water dynamics than that of pure osmotic cell swelling, emphasizing the need to focus on the multifaceted cellular actions of K^+^. Such biophysical insights are expected to not only advance our understanding of basic cell physiology but also suggest potential endogenous MRI contrast mechanisms for detecting specific K^+^- driven cellular events.

## Materials and Methods

### Cell preparation

We used non-excitable T-lymphocytes cells (or Jurkat), which have a spherical shape suitable for modeling of cell structure, are stable in terms of membrane potential.

Jurkat cells were cultured in Roswell Park Memorial Institute (RPMI) 1640 medium with 10 (v/v) fetal bovine serum (FBS, Thermo Fisher Scientific, Waltham, MA, USA) and 1 (v/v) penicillin/streptomycin (Thermo Fisher Scientific, Waltham, MA, USA). The composition of the cell medium during MR scanning was: 4.2 mM KCl, 145.8 mM NaCl, 20 mM 4-(2-hydroxyethyl)-1-piperazineethanesulfonic acid (HEPES, Thermo Fisher Scientific, Waltham, MA, USA), 4.5g/L glucose (Thermo Fisher Scientific, Waltham, MA, USA), and 10μM ethylenediaminetetraacetic acid (EDTA, Thermo Fisher Scientific, Waltham, MA, USA) with pH 7.2.

To compare the osmotic changes induced by sodium ions (Na^+^) and the membrane potential changes induced by potassium ions (K^+^), the cells were placed in media with manipulated concentrations of each ion. We prepared three different media containing additional NaCl of -5, -10, and -15 mM, which can derive resulting osmolarity of -10, - 20, and -30 mM, to control the osmotic pressure of each medium. We also prepared three additional media with [K^+^] of 20, 40, and 80 mM to control the membrane potential of cells within each medium. For MRI measurements, each cell suspension (prepared to yield sufficient signal for the requested slice thickness) was placed into each cylindrical well perforated into a double-sided spherical acrylic phantom (Figure 1a) and centrifuged (1,100 RPM, 5 min) to form a pellet as illustrated in Figure 1b.

### MRI data acquisition

The acrylic phantom with cell pellets was scanned on a 9.4 T animal MRI scanner (BioSpin, Bruker). All data were collected in the form of 1D images with varying echo time (TE) using scan parameters such as spatial resolution 0.25 mm, number of pixels 128, and slice thickness 1 mm (Figure 1c). For T2 estimation, MRI signals were acquired using a single spin-echo sequence with a repetition time (TR) of 2000 ms and 40 TEs ranging from 8.1 ms to 85.4 ms on a logarithmic scale. For MT estimation, MRI signals were acquired using a selective inversion recovery multi-echo spin-echo sequence (IR-MESE) with 40 inversion times (TI) ranging from 0.24 ms to 10000 ms on a logarithmic scale and 16 TEs increasing linearly from 8.1 ms to 129.6 ms which can be acquired multiple data points along the T2 relaxation decay curve, thus enabling more precise estimation of MT parameters. To ensure sufficient recovery of longitudinal magnetization, TR was set to 15000ms.

### Data analysis

Assuming that there are two hydrogen proton pools: free water pool and bound pool, the dynamics between two pools can be expressed with their own longitudinal relaxation rates (𝑅_1*f*_ for free water pool and 𝑅_1𝑏_ for bound pool) and rates of magnetization transfer (𝑘_𝑏*f*_, bound pool è free water pool; 𝑘_*f*𝑏_, free water pool è bound pool). Assuming that *M*_*f*_ 𝑡 and *M*_𝑏_ 𝑡 represent the longitudinal magnetizations of the free water pool and bound pool, respectively, *M*_*f*_ 𝑡 was given by

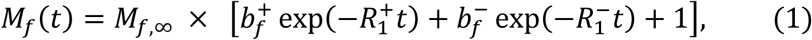

where *M*_*f*,∞_ is the longitudinal magnetization at equilibrium state, and R1^±^ and bf^±^ are defined as

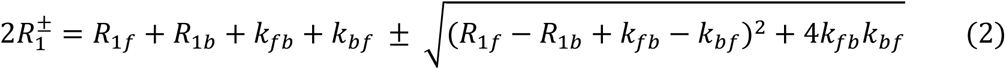

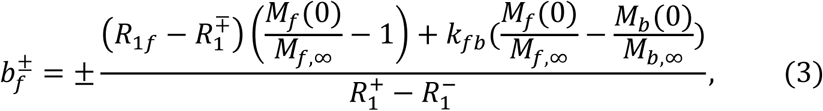

The ratio *M*_𝑏_ /*M*_𝑏,∞_ was calculated using the Bloch equations, based on the duration of the inversion pulse of the IR-MESE sequence, as demonstrated in previous studies^50,51^. The resulting MRI signal acquired with 16 TEs was fitted with mono-exponential decay to yield *M*_*f*_ 𝑡 . To avoid stimulated echo contamination inherent to MESE sequences, only even numbered TEs were used for fitting^52^. *M*_*f*_ 𝑡 was fitted to yield 𝑏_f_^±^, 𝑅_1_^±^, and *M*_*f*,∞_ using 40 inversion times (TI) according to Equation (1). This fitting was performed using a particle swarm optimization algorithm^53^ in MATLAB (R2022b, MathWorks Inc., Natick, MA, USA). PSR was estimated by the ratio of the transfer rates 𝑘_*f*𝑏_ and 𝑘_𝑏*f*_ according to the principle of microscopic reversibility^50,51,54^. Furthermore, assuming that R1f and R1b are equal, the equation for 𝑘_*f*𝑏_/𝑘_𝑏*f*_ simplifies to

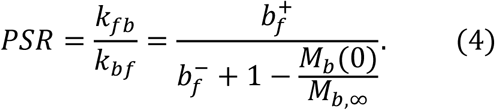

The detailed method for determining this value was described in previous MT _studies_50,51,55.

### Cell volume measurement

To measure cell volume using a microscope, all cells were placed in petri dishes with prepared media, where the osmolarity and [K^+^] were controlled. Digital images were captured using an Olympus CKX53 microscope at 20x magnification (Figure 1d). The radii of over 1000 cells from the images by medium were analyzed using the “imfindcircles” function, which employs the Circular Hough Transform algorithm to detect circles in MATLAB. Cell volume distributions were plotted assuming spherical shape of the cells (Figure 2). The mean cell volume 〈*V*_*c*_〉 was estimated using the cell volume distribution.

### Modeling

We assumed that Jurkat cells with a spherical shape would form a close-packed structure as a result of centrifugation (Figure 1b). In this structure, the intracellular space fraction within a voxel (*v*_*i*_) is maintained by reducing the number of cells as the cell volume increases with the expansion of the cell radius^29,30^. For example, the *v*_*i*_ of the FCC (Face Centered Cubic) structure, which has the most effective packing structure, is about 74 . *v*_*i*_ can be estimated by the product of the cell number density (*ρ*_*cell*_) and the average cell volume (〈*V*_*c*_〉)^56^.

In a similar mechanism to represent *v*_*i*_ using *ρ*_*cell*_, the longitudinal magnetization of bound pool at equilibrium state (*M*_𝑏,∞_) can be estimated by the product of *ρ*_*cell*_ and the mean longitudinal magnetization of bound pool per cell (〈*M*_𝑏,*c*_〉), and by definition, *M*_𝑏,∞_ is given by the product of the pool size ratio (PSR) and the magnetization of free water pool (*M*_*f*,∞_). Therefore, 〈*M*_𝑏,*c*_〉 can be expressed as

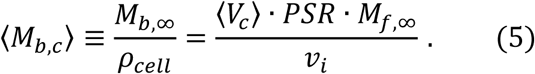

## Acknowledgement

This work was supported by the Brain Research Program through the National Research Foundation of Korea (NRF) funded by the Ministry of Science, ICT & Future Planning (NRF-2023R1A2C3007075).

## References

1. Macknight A, Leaf A. Regulation of cellular volume. Physiological reviews 1977;57(3):510–573

2. Adrian R. The effect of internal and external potassium concentration on the membrane potential of frog muscle. The Journal of Physiology 1956;133(3):631

3. Udensi UK, Tchounwou PB. Potassium homeostasis, oxidative stress, and human disease. International journal of clinical and experimental physiology 2017;4(3):111

4. Yu SP. Regulation and critical role of potassium homeostasis in apoptosis. Progress in neurobiology 2003;70(4):363–386

5. Ramsey CL. Biologically Structured Water (BSW)-A Review (Part 1): Structured Water (SW) Properties, BSW and Redox Biology, BSW and Bioenergetics. Journal of Basic & Applied Sciences 2023;19(174-201

6. Higgins MJ, Polcik M, Fukuma T, et al. Structured water layers adjacent to biological membranes. Biophysical journal 2006;91(7):2532–2542

7. Privalov PL, Crane-Robinson C. Role of water in the formation of macromolecular structures. European biophysics journal 2017;46(3):203–224

8. Tros M, Zheng L, Hunger J, et al. Picosecond orientational dynamics of water in living cells. Nature Communications 2017;8(1):904

9. Fayer MD, Levinger NE. Analysis of water in confined geometries and at interfaces. Annual review of analytical chemistry 2010;3(89-107

10. Persson E, Halle B. Cell water dynamics on multiple time scales. Proceedings of the National Academy of Sciences 2008;105(17):6266–6271

11. Disalvo EA, Lairion F, Martini F, et al. Structural and functional properties of hydration and confined water in membrane interfaces. Biochimica et Biophysica Acta (BBA)-Biomembranes 2008;1778(12):2655–2670

12. Dass AV, Jaber M, Brack A, et al. Potential role of inorganic confined environments in prebiotic phosphorylation. Life 2018;8(1):7

13. Watson JL, Seinkmane E, Styles CT, et al. Macromolecular condensation buffers intracellular water potential. Nature 2023;623(7988):842-852

14. Lang X, Shi L, Zhao Z, et al. Probing the structure of water in individual living cells. Nature Communications 2024;15(1):5271

15. Orlikowska-Rzeznik H, Versluis J, Bakker HJ, et al. Cholesterol Changes Interfacial Water Alignment in Model Cell Membranes. Journal of the American Chemical Society 2024;146(19):13151–13162

16. Tarun OB, Hannesschläger C, Pohl P, et al. Label-free and charge-sensitive dynamic imaging of lipid membrane hydration on millisecond time scales. Proceedings of the National Academy of Sciences 2018;115(16):4081–4086

17. Shi L, Hu F, Min W. Optical mapping of biological water in single live cells by stimulated Raman excited fluorescence microscopy. Nature communications 2019;10(1):4764

18. Sterpone F, Stirnemann G, Laage D. Magnitude and molecular origin of water slowdown next to a protein. Journal of the American Chemical Society 2012;134(9):4116–4119

19. Didier M, Tarun O, Jourdain P, et al. Membrane water for probing neuronal membrane potentials and ionic fluxes at the single cell level. Nature communications 2018;9(1):5287

20. de Coene Y, Jooken S, Deschaume O, et al. Label Free Imaging of Membrane Potentials by Intramembrane Field Modulation, Assessed by Second Harmonic Generation Microscopy. Small 2022;18(18):2200205

21. Henkelman RM, Huang X, Xiang QS, et al. Quantitative interpretation of magnetization transfer. Magnetic resonance in medicine 1993;29(6):759–766

22. Sled JG, Pike GB. Quantitative imaging of magnetization transfer exchange and relaxation properties in vivo using MRI. Magnetic Resonance in Medicine: An Official Journal of the International Society for Magnetic Resonance in Medicine 2001;46(5):923–931

23. Min K, Chung S, Lee S-K, et al. Responses to membrane potential-modulating ionic solutions measured by magnetic resonance imaging of cultured cells and in vivo rat cortex. eLife Sciences Publications, Ltd: 2025.

24. Sehy JV, Ackerman JJ, Neil JJ. Water and lipid MRI of the Xenopus oocyte. Magnetic Resonance in Medicine: An Official Journal of the International Society for Magnetic Resonance in Medicine 2001;46(5):900–906

25. Sehy JV, Ackerman JJ, Neil JJ. Evidence that both fast and slow water ADC components arise from intracellular space. Magnetic Resonance in Medicine: An Official Journal of the International Society for Magnetic Resonance in Medicine 2002;48(5):765–770

26. Harkins KD, Galons JP, Secomb TW, et al. Assessment of the effects of cellular tissue properties on ADC measurements by numerical simulation of water diffusion. Magnetic Resonance in Medicine: An Official Journal of the International Society for Magnetic Resonance in Medicine 2009;62(6):1414–1422

27. Henkelman R, Stanisz G, Graham S. Magnetization transfer in MRI: a review. NMR in Biomedicine: An International Journal Devoted to the Development and Application of Magnetic Resonance In Vivo 2001;14(2):57–64

28. Stanisz GJ, Kecojevic A, Bronskill M, et al. Characterizing white matter with magnetization transfer and T2. Magnetic Resonance in Medicine: An Official Journal of the International Society for Magnetic Resonance in Medicine 1999;42(6):1128–1136

29. Rintoul M, Torquato S. Computer simulations of dense hard sphere systems. The Journal of chemical physics 1996;105(20):9258–9265

30. Farr RS, Groot RD. Close packing density of polydisperse hard spheres. The Journal of chemical physics 2009;131(24):

31. Jentsch TJ. VRACs and other ion channels and transporters in the regulation of cell volume and beyond. Nature Reviews Molecular Cell Biology 2016;17(5):293

32. Völkl H, Lang F. Effect of potassium on cell volume regulation in renal straight proximal tubules. The Journal of membrane biology 1990;117(113-122

33. Delpire E, Gagnon KB. Water homeostasis and cell volume maintenance and regulation. Current topics in membranes 2018;81(3-52

34. Cheung RK, Grinstein S, Dosch HM, et al. Volume regulation by human lymphocytes: characterization of the ionic basis for regulatory volume decrease. Journal of Cellular Physiology 1982;112(2):189–196

35. Bui A, Wiley J. Cation fluxes and volume regulation by human lymphocytes. Journal of Cellular Physiology 1981;108(1):47–54

36. Jara H, Sakai O, Farrher E, et al. Primary multiparametric quantitative brain MRI: state-of-the-art relaxometric and proton density mapping techniques. Radiology 2022;305(1):5–18

37. Alexander AL, Hurley SA, Samsonov AA, et al. Characterization of cerebral white matter properties using quantitative magnetic resonance imaging stains. Brain connectivity 2011;1(6):423–446

38. Birkl C, Doucette J, Fan M, et al. Myelin water imaging depends on white matter fiber orientation in the human brain. Magnetic resonance in medicine 2021;85(4):2221–2231

39. MacKay AL, Laule C. Magnetic resonance of myelin water: an in vivo marker for myelin. Brain plasticity 2016;2(1):71–91

40. Min K, Chung S, Lee S-K, et al. Detection of changes in membrane potential by magnetic resonance imaging. bioRxiv 2024;2024.04. 02.587661

41. Abe Y, Van Nguyen K, Tsurugizawa T, et al. Modulation of water diffusion by activation-induced neural cell swelling in Aplysia Californica. Scientific reports 2017;7(1):6178

42. Le Bihan D. The ‘wet mind’: water and functional neuroimaging. Physics in Medicine & Biology 2007;52(7):R57

43. Le Bihan D. Diffusion MRI: what water tells us about the brain. EMBO molecular medicine 2014;6(5):569–573

44. Dombeck DA, Blanchard-Desce M, Webb WW. Optical recording of action potentials with second-harmonic generation microscopy. Journal of Neuroscience 2004;24(4):999–1003

45. Lee HJ, Jiang Y, Cheng J-X. Label-free optical imaging of membrane potential. Current opinion in biomedical engineering 2019;12(118-125

46. Wang Y-H, Zheng S, Yang W-M, et al. In situ Raman spectroscopy reveals the structure and dissociation of interfacial water. Nature 2021;600(7887):81-85

47. Grossman RI. Magnetization transfer in multiple sclerosis. Annals of Neurology: Official Journal of the American Neurological Association and the Child Neurology Society 1994;36(S1):S97–S99

48. Davoudi Y, Foroughipour M, Torabi R, et al. Diffusion weighted imaging in acute attacks of multiple sclerosis. Iranian Journal of Radiology 2016;13(2):e21740

49. Pasco A, Minassian AT, Chapon C, et al. Dynamics of cerebral edema and the apparent diffusion coefficient of water changes in patients with severe traumatic brain injury. A prospective MRI study. European radiology 2006;16(1501-1508

50. Li K, Zu Z, Xu J, et al. Optimized inversion recovery sequences for quantitative T1 and magnetization transfer imaging. Magnetic Resonance in Medicine 2010;64(2):491–500

51. Gochberg DF, Kennan RP, Robson MD, et al. Quantitative imaging of magnetization transfer using multiple selective pulses. Magnetic Resonance in Medicine: An Official Journal of the International Society for Magnetic Resonance in Medicine 1999;41(5):1065–1072

52. McPhee KC, Wilman AH. Limitations of skipping echoes for exponential T2 fitting. Journal of Magnetic Resonance Imaging 2018;48(5):1432–1440

53. Kennedy J, Eberhart R. Particle swarm optimization. ieee: 1995.

54. Trujillo P, Summers PE, Smith AK, et al. Pool size ratio of the substantia nigra in Parkinson’s disease derived from two different quantitative magnetization transfer approaches. Neuroradiology 2017;59(1251-1263

55. Gochberg DF, Gore JC. Quantitative imaging of magnetization transfer using an inversion recovery sequence. Magnetic Resonance in Medicine: An Official Journal of the International Society for Magnetic Resonance in Medicine 2003;49(3):501–505

56. Springer Jr CS, Baker EM, Li X, et al. Metabolic activity diffusion imaging (MADI): I. Metabolic, cytometric modeling and simulations. NMR in Biomedicine 2023;36(1):e4781

