## Supplementary materials for "Exploring Cellular Water Dynamics associated with Potassium Ion Changes Using Magnetic Resonance Imaging"

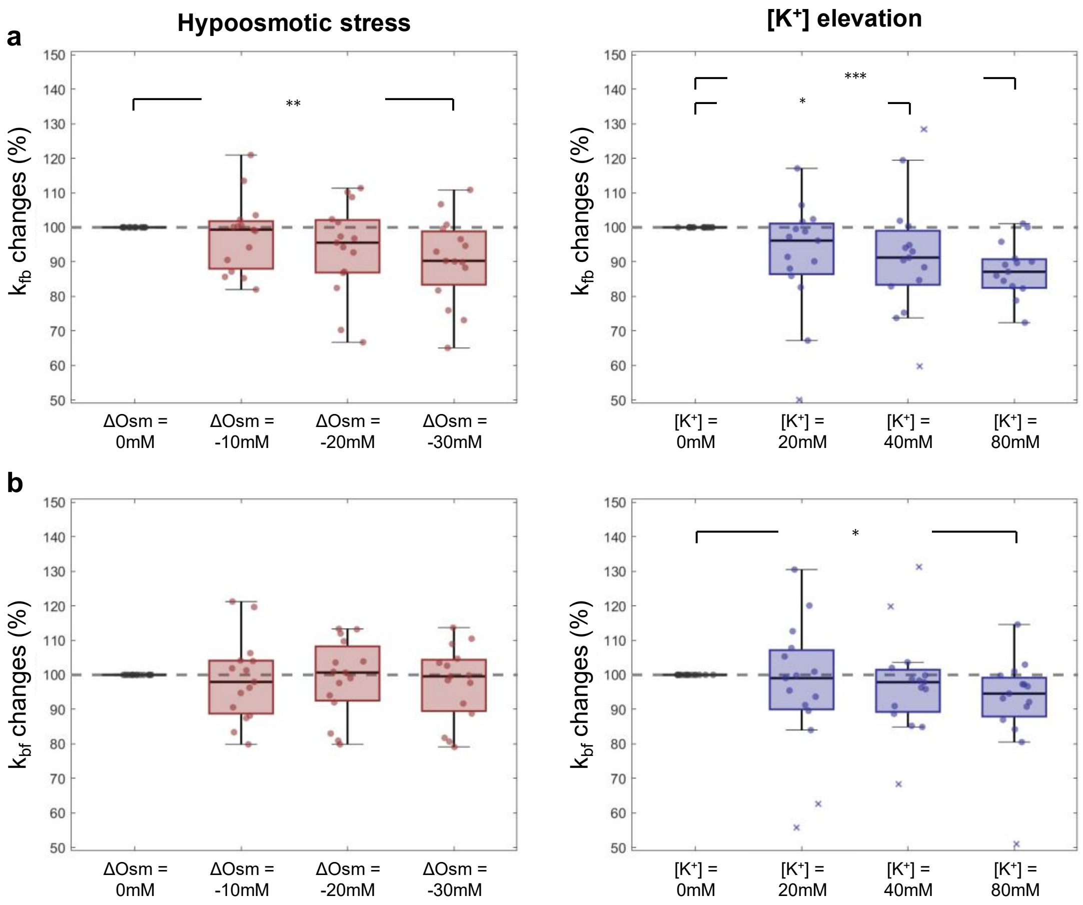


**Supplementary Figure 1**. Estimation of magnetization transfer rates, $k_{fb}$ and $k_{bf}$ ($k_{fb}$: the transfer rate from the free water pool to the bound pool; $k_{bf}$: the transfer rate in the opposite direction). **a.** $k_{fb}$ changes under hypoosmotic stress (three on the left) and [K$^+$] elevation (three on the right) conditions. $k_{fb}$ was reduced in both cases, which is consistent with a previous study that measured transfer rates at different protein concentrations[53]. **b.** $k_{bf}$ changes under hypoosmotic stress (three on the left) and [K$^+$] elevation (three on the right) conditions. Unlike the previous study mentioned above[53] where no change in $k_{bf}$ was reported despite changes in PSR and $k_{fb}$, in our study $k_{bf}$ decreased under high [K$^+$] conditions. Statistical significance was marked with asterisks (*: p < 0.05; **: p < 0.01; ***: p < 0.001).

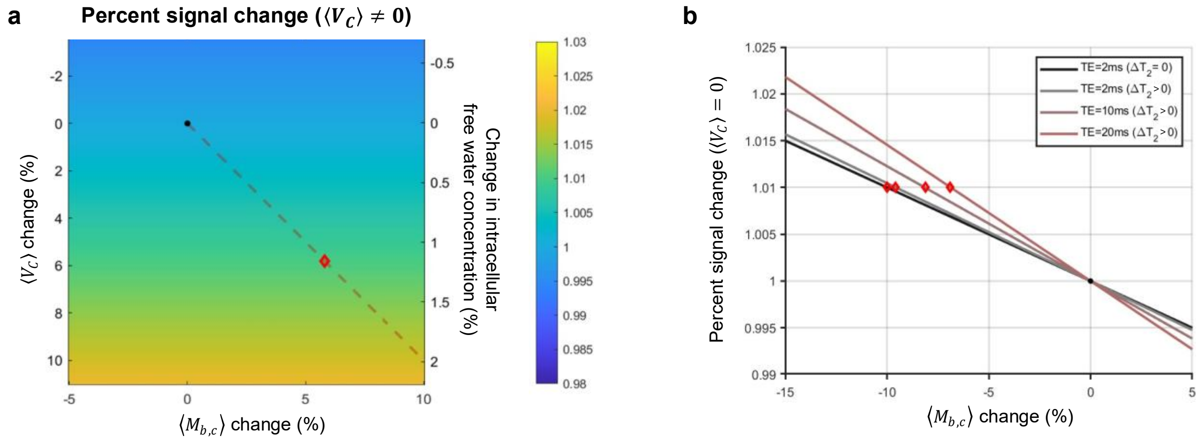


**Supplementary Figure 2**. Simulation of percent signal change induced by changes in water dynamics. A spoiled gradient-echo sequence was used simulation. Sufficient dummy scans were applied before the main sequence to ensure that the longitudinal magnetization ($M_z$) reached steady state. Simulation parameters were as follows: $T_1$ = 1 s for water, $T_2$ = 70 ms for free water pool, $T_2$ = 0.1 ms for bound pool, TR/TE = 5 ms/2 ms (**a**), TR/TE = 5 ms/2ms, 50 ms/10 and 20 ms (**b**), flip angle = 4.6°, PSR = 10%, and intracellular fraction = 82%. **a.** Percent signal change in the presence of cell swelling ($\Delta\langle V_c \rangle \neq 0$). Since $\langle V_c \rangle$ and $M_{f,\infty}$ a have linear relationship in Figure 4b, i.e., $\Delta\langle V_c \rangle \approx 5\Delta M_{f,\infty}$, the change in $\langle V_c \rangle$ (left vertical axis) and the change in intracellular free water concentration (right vertical axis) can be considered as equivalent y-axis. The red dashed line reflects the relationship between the changes in $\langle M_{b,c} \rangle$ and $\langle V_c \rangle$ shown in Figure 5c. The red diamond indicates a 1% signal change calculated for a 5.8% increase in average cell volume ($\langle V_c \rangle$), considering both $M_{f,\infty}$ and $\langle M_{b,c} \rangle$. **b.** Percent signal change due to redistribution of bound water pool without cell swelling. Simulation results showed that the increase in percent signal arises from a transition of part of the bound water pool with short $T_2$ to the free water pool with long $T_2$.The increase in $T_2$ shown in Figure 4c was considered as the cases of $\Delta T_2 > 0$ compared to the control case of $\Delta T_2 = 0$. To better demonstrate the percent signal increase due to $T_2$ increase, three increasing TEs (= 2, 10, 20 ms) were assumed to enhance the $T_2$ signal contrast. There was no significant difference according to the change in TR (≤ 50ms). The 1% signal increase was marked with a red diamond. For the case of $\Delta T_2 > 0$ with TE = 20 ms, a 6.9% decrease in the bound pool ($\langle M_{b,c} \rangle$) resulted in a 1% signal increase. ($\langle V_c \rangle$: the average cell volume, $M_{f,\infty}$: the longitudinal magnetization of free water pool at equilibrium state, $\langle M_{b,c} \rangle$: the average longitudinal magnetization of bound pool per cell).
